# Understanding robustness and enabling remediation of neuroimaging-based deep neural networks for digital security: Semantic segmentation of brain tumors as illustrative study

**DOI:** 10.64898/2026.11.10.765546

**Authors:** Mayur Mankar, Bhaskar Roy, Prasun K. Roy

**Affiliations:** Neuroimaging Laboratory, Life Sciences Dept, Shiv Nadar University (Institution of Eminence), Delhi NCR, 201314, India; Dept. of Data Science and Engineering, Indian Institute of Science Education and Research, Bhopal, 462066, India; SNU-Dassault Centre of Excellence for Research and Innovation, Shiv Nadar University, Greater Noida, Delhi NCR, 201314

**Keywords:** Cybersecurity, Neuro-oncology, Adversarial attacks, Brain tumor, Neural network

## Abstract

**Background:** Applications of deep learning models in neuroimaging have significantly advanced translational neuroscience, offering substantial benefits to human health. However, recent cybersecurity threats have undermined the reliability of digital neuroscience tools, posing challenges to the clinical neuroscience community. A critical issue is the automated semantic segmentation of malignant or premalignant brain tumors, essential for neurosurgery and radiotherapy planning.

**New methods:** This study investigates the robustness of customary deep learning models such as Convolutional Neural Networks (CNNs) and more recent Vision Transformers (ViTs) in semantic segmentation against adversarial attacks. We assess the performance of CNNs and ViTs under ideal nonhostile conditions and then under cybersecurity threats.

**Results:** Our findings reveal that while CNNs perform well in ideal conditions, their efficiency sharply declines under adversarial perturbations. In contrast, ViTs maintain their segmentation performance even when subjected to adversarial attacks. This study is the first to apply Visual Transformers for cybersecurity challenge, leveraging advanced capabilities of VITs to enhance robustness against digital perturbations.

**Comparison with existing methods:** Compared to CNNs, which are widely used for neuroimages but vulnerable to adversarial attacks, ViTs demonstrate superior resilience. Our research highlights that ViTs are as effective as CNNs under ideal conditions but offer enhanced protection from cybersecurity angle, making them a more robust choice for neuroimaging applications.

**Conclusion:** Our results indicate that for large-scale segmentation operation in neuroimaging, such as lesion segmentation tasks, ViT-based models should be preferred over CNN-based models. This shift could provide inherent protection against prevalent digital threats, ensuring more reliable and secure neuroimaging diagnostics.

## 1. INTRODUCTION

Digital health methodologies deep neural networks have become well-utilized for neuroimaging analysis tasks like segmentation of regions of interest, as brain lesions, atrophy or tumor detection, with utility for diagnosis, prognostics, grading and therapy planning. However, recent hazardous experiences, from the angle of cybersecurity risks, do demonstrate that carefully-engineered adversarial attacks can jeopardize neuroinformatics and neuroradiological procedures and seriously compromise the performance of imaging-based deep learning systems under the deliberate effect of subtle and hostile perturbations. This raises security concerns about the use of these computational methods of neuroimaging in clinical diagnostic settings. Our study is the first of its kind to investigate the robustness of deep learning-based MRI diagnostic systems using adversarial intrusions.

From the perspective of medical neuroscience, a most acute clinical issue is brain tumors, as it is the largest cause of malignant mortality in children and youth, and the prevalence globally is slated to be about 1,000,000 cases by the end of the next decade. In fact, the 5-year survival rate for such a tumor, like glioblastoma, is only 1 to 2 years. Neuroimaging analysis of patients plays a crucial role in diagnosing neurological disorders as well as brain tumors. Early detection of such disorders can be vital for improving treatment outcomes. Magnetic Resonance Imaging (MRI), a common non-invasive strategy, has been used to diagnose lesions in the brain. However, manually analysing these structural MRI images is a tiring and time-consuming which, till now is only successfully done by professional neuroradiologists whose numbers and availability are tightly constrained worldwide, highlighting the serious need for automatic and robust brain tumor segmentation systems. Deep neural networks (DNNs) have emerged as effective tools for automating brain image segmentation tasks. Popular DNNs for neuroimaging applications include CNNs and ViTs. The utility of transformer architectures has impressively come to the public awareness after the development of ChatGPT, using generative pre-training transformers (GPT).

However, automated segmentation systems are vulnerable to adversarial cyberattacks, which is a cause for great concern. Considering the negative impacts of such attacks, our paper analyzes the robustness of DNNs against adversarial attacks in white-box adversarial attack settings. We highlight the XAI (explainable artificial intelligence) aspect by applying XAI tools such as Grad-CAM [1] and approach the matter both from a model-based and data-based perspective, and show the high robustness of security-effectivity of the procedure that we have developed. The potentiality of our research can have considerable impact, showing that instead of the current stress of using customary convolutional neural network method, the medical imaging diagnostic industry may consider giving high emphasis of the more recent digital discoveries, such as transformers. We would also like to underscore the substantial novelty in our work since, as far as we are aware, application to the neuroimaging and medical imaging scenario has not been done earlier regarding the cybersecurity and robustness angle of the segmentation process for pathological lesions.

## 2. MATERIAL AND METHODS

### 2.1. MRI Scanning Acquisition

The MRI scans of patients with brain tumors [2] were acquired at the General Hospital, Tianjin Medical University, China, and also from Nanfang Hospital, Guangzhou, China between the 2005 to 2010. The scan-set has 3064 T1w brain CE-MRI from 233 patients. The image resolution was set as 512 × 512 and the pixel size was set to 0.49 × 0.49 *mm*^2^. Institutional human ethics approval was there. These MRI images were from patients having one of the three prevalent types of primary brain tumors, namely meningioma (708 patients), glioma (1426 patients) and pituitary tumor (930 patients). Tumor class labels and the tumor boundary were manually defined and marked by three experienced neuroradiologists in the same hospitals (figure1).

### 2.2. Deep Neural Networks

We test a plethora of models from each architecture family to better understand and differentiate between ViTs and CNNs. We employ models that differ in terms of input patch-size, number of variables, and datasets on which they were pre-trained.

#### 2.2.1. Convolutional Neural Networks

Recently there has been an interest in utilizing U-Net models in analysing neuroimages. It is a classic image–to–image translator model networks with an encoder-decoder architecture and is widely used in brain image segmentation. U-Net++, a version of U-Net has been developed as an improved U-Net by having an architecture that uses dense skip and nested connections. Each network architecture comes with its own advantages in the use of the encoder and decoder. We have done analysis on U-Net [3], U-Net++ [4], Dynamic-UNet [5], Dense-UNet [6] and Res-UNet [7] due to their success in analysing neuroimages.

#### 2.2.2. Vision Transformers

ViTs [8] are derived from the transformer architecture for the use of analysing images, revolutionizing the area of computer vision. By converting the input image pixels into latent representations and using a self-attention mechanism to find long-range dependencies and interactions between the image parts, ViT can recognize the intrinsic patterns that exist in images. This model works by breaking down the input image into non-overlapping fixed-size patches and turning each patch into a representation, which are called patch embeddings. The transformer encoder receives these patches embedding as input after having undergone linear transformation. Patches allow the model to view the whole image which captures the global context. To encode positional information positional embeddings are added to the patch embeddings, this helps the model maintain awareness of the spatial arrangement of the patches in the original image. We dive into more particulars about ViTs in the upcoming sections.

At the core of the ViT architecture is the transformer encoder. It consists of several layers of multiheaded self-attention (MSA) and Multi-Layer Perceptrons (feed-forward neural networks). The multiheaded self-attention (MSA) heads make it possible for the model to detect local and global patch relationships, and non-linearity is added by feed-forward networks to enhance the features. The output of the final encoder is usually pooled, and a classification head is used for making predictions. To obtain image masks for our image segmentation purposes a specialized decoder block is utilized. We have done analysis on Swin-UNet [9] and Trans-UNet [10].

### 2.3. Semantic Segmentation

Currently, CNN-based models have seen great success in the use for medical image segmentation due to its powerful representation ability. With the advancement of deep CNN, U-Net is proposed in medical image analysis. Because of the simple structure and better performance of the U-shaped structure, various U-Net models are constantly developed and produced, such as U-Net++, Dynamic-UNet, Dense-UNet and Res-UNet.

Vision Transformers are an alternative to traditional Convolutional Neural Networks (CNNs) for brain image segmentation tasks. One drawback of ViT compared to CNN is that it requires its own large dataset to be pre-trained on. Recently, several excellent works have been done based on ViT. Swin-UNet and Trans-UNet are two cutting-edge algorithms proposed for medical image segmentation. Swin-UNet is entirely transformer base. Trans-UNet is a combination of transformer and CNN model. And is made up of a transformer encoder and a convolutional decoder.

### 2.4. Adversarial Attack and perturbations

Adversarial attacks involve unnoticeable manipulations of training data or generation of new manipulated training data with the intention of causing algorithmic errors. The perturbations created in these attacks are often imperceptible to humans, necessitating the need for algorithms to detect them. Crafting these attacks requires meticulous calculations to exploit a particular algorithm or model’s vulnerabilities. Adversarial attacks are further classified into white-box and black-box attacks, depending on whether the model architecture, training data, and other relevant information are available to the attacker or not.

Adversarial attacks on brain image segmentation models involve generating adversarial examples that deceive the model into predicting incorrect segmentation masks. In our experiments, we aim to apply such adversarial attacks to models trained for brain tumor image segmentation. By presenting these adversarial images during the evaluation phase, we observe that the models produce erroneous segmentation masks, highlighting the susceptibility of brain image segmentation models to adversarial manipulation.

## 3. THEORY AND CALCULATION

### 3.1. Brain Image Semantic Segmentation

In this paper, we attend to brain image segmentation task with the use of DNNs. For a binary classification problem, given a dataset {(*x*_*i*_, *y*_*i*_)}_*i*=1,…,*N*_where *x*_*i*_ ∈ ℝ^*d*^ represents an input image and *y*_i_ ∈ {0,1} represents the binary label (0 for background, 1 for object) (figure 2), a DNN classifier *h* with parameters *θ* predicts the class of each pixel in the input image *x*_i_: *h* (*x*_*i*_) = argmax_*c*=0,1_*p*_*c*_(*x*_*i*_, *θ*), where

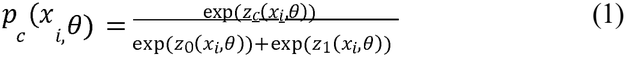

and *z*_*c*_(*x*_i_,*θ*) is the logits output of the network for class *c* (where *c* can be either 0 or 1). The probability *p*_*c*_(*x*_*i*,_ *θ*) represents the likelihood of each pixel in *x*_*i*_ that belongs to class *c*.

**Figure 1.**
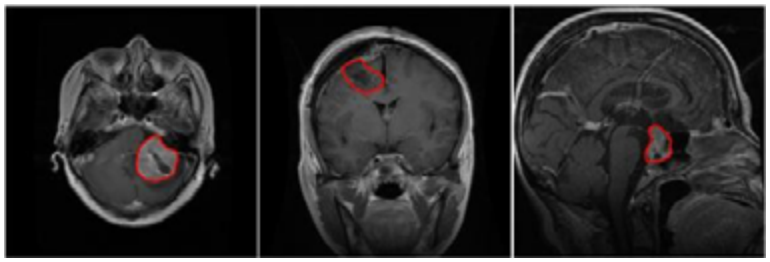
Illustration of three typical brain tumors (1) meningioma (2) glioma (3) pituitary tumor as accessed from the dataset.

**Figure 2.**
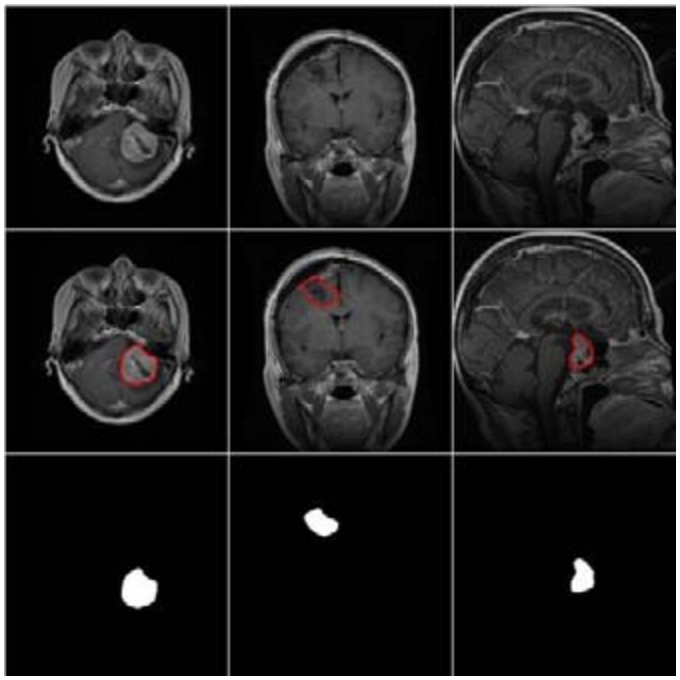
Binary semantic segmentation of brain MRI image is done to locate the tumor, where tumor is given white color and remaining part is given black color.

Back-propagation is used to update the model parameters *θ*, to minimize the classification loss, which is typically the binary cross-entropy loss:

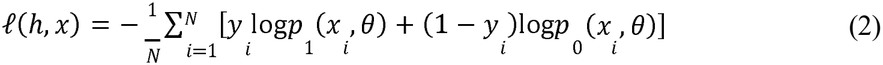

Here, *l*(*h,x*) measures the discrepancy between the predicted probabilities of assigned label and the true binary labels, and the objective is to minimize this loss to improve the model’s performance in segmenting the objects of interest from the background. N denotes the total number of image-label pairs in the dataset.

Given a pretrained DNN model *h* and an input image *x* with its associated binary segmentation label *y*, an adversarial attack aims to maximize the segmentation error of the DNN model while ensuring that the adversarial example *xadv* remains within a small *E*-ball centered around the original image *x*, i.e., ‖*xadv−x*‖*p* ≤*∈*. Here ‖·‖*p* denotes the *Lp*-norm, with *p* = ∞ being the most commonly used as it is consistent with human perception.

### 3.2. Adversarial Attacks

Adversarial attacks in binary semantic segmentation can be categorized into two, targeted or untargeted. In a targeted attack, the goal of the attacker is to create an adversarial example *xadv* that causes the DNN to misclassify the segmentation mask to a specific target class (e.g., background if the true class is object). In an untargeted attack, the objective of the attacker is to find an adversarial example *xadv* that results in any misclassification, irrespective of the specific erroneous class.

These attacks can either be created in a white-box or black-box environment. In a white-box context, the gradients of the loss function with respect to the input, taken straight from the target model, are used to create the adversarial instances. In a black-box setting, attacks are executed using a surrogate model or estimated adversarial gradients. In this paper, we have focused on untargeted attacks in the white-box setting under the *L*_∞_ perturbation constraint. To create adversarial instances for untargeted white-box attacks, the following constrained optimization problem is solved:

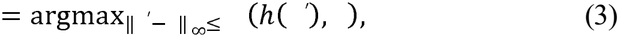

where (·) is the segmentation loss obtained from (2), and is the true binary label. This formulation aims to maximize the classification loss, thereby leading to significant errors in the segmentation output, while ensuring the perturbations are within the specified bound. A variety of methods have been developed to produce such adversarial examples, and we will go over a selection of most distinctive and cutting-edge attacks. The model is first trained on normal dataset and a perturbed image (with adversarial attack) is fed to the model to test its analysis capabilities (Figure 3).

**Figure 3.**
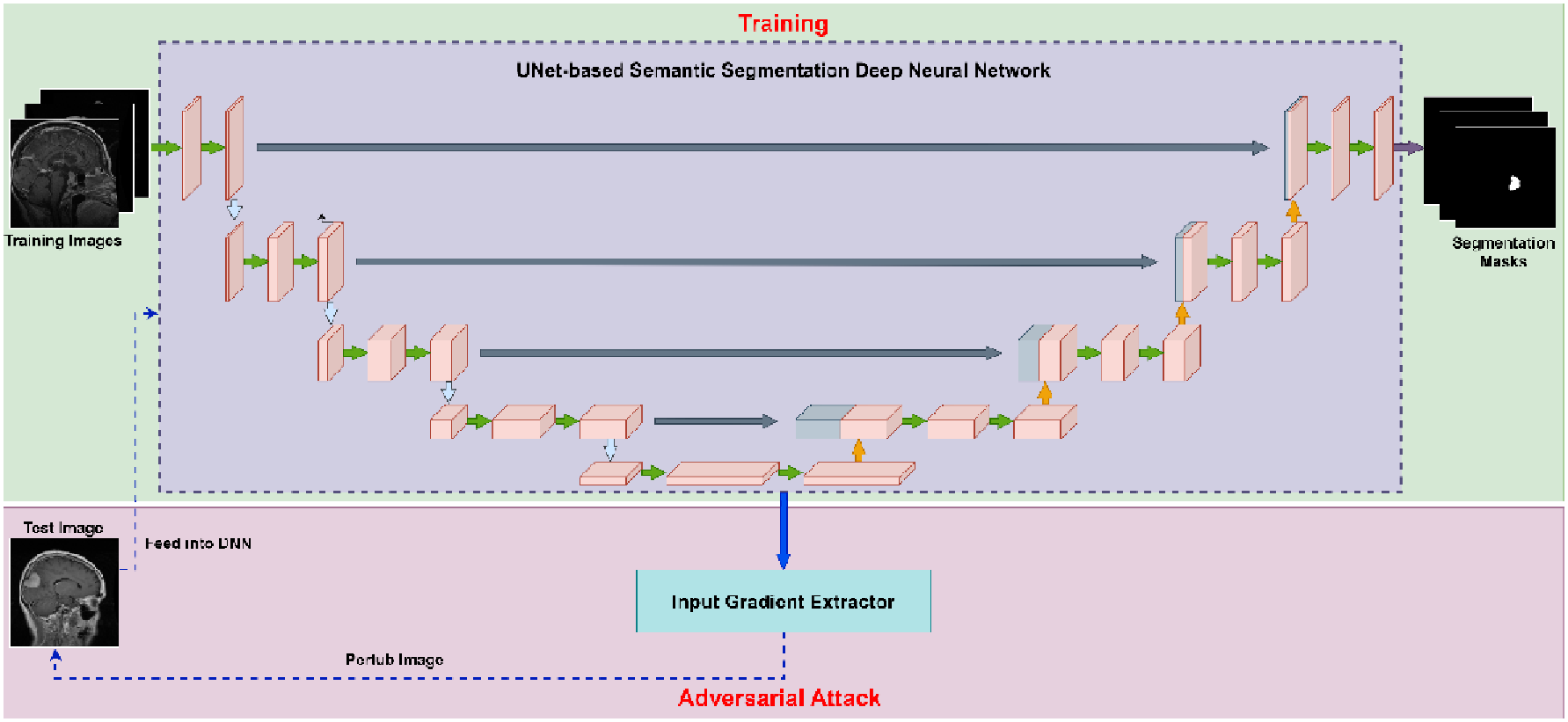
The schema representing the pipeline of training Deep Neural Networks (DNNs) and generating adversarial perturbations.

#### 3.2.1. Fast Gradient Sign Method (FGSM)

FGSM [11] alters standard examples by along the input gradient direction for one step:

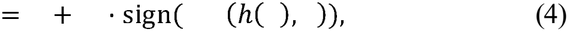

Where, sign(·) is the function returning the sign of the input ,(·) is the loss function obtained from (2), and (·) represents the gradient of (·) w.r.t x.

#### 3.2.2. Basic Iterative Method (BIM)

An iterative version of FGSM is BIM [12]. Different to FGSM, BIM iteratively perturbs the input image with a smaller step size:

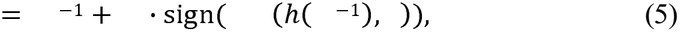

Where, *x*^*t*^is the adversarial example at the *t*-th step (*x*^0^ = *x*) and, *α* is the step size. Usually, the step size is set to *ϵ*/*T* ≤ *α* < *ϵ* for a total *T* steps of perturbation.

#### 3.2.3. Projected Gradient Descent (PGD)

PGD [13] perturbs a standard example *x* for *T* steps with smaller step size. PGD projects the adversarial example back onto the *ϵ*-ball of *x* after each step if it exceeds:

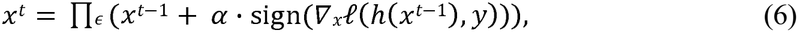

Where *x*^*t*^ is the adversarial example at the *t*-th step (*x*^0^ = *x*), *α* is the step size, ∏(·) is the projection function. PGD uses random start for *x*^0^ = *x* + *U*^*d*^(−*ϵ, ϵ*), where *U*^*d*^(−*ϵ, ϵ*) is the uniform distribution between −*ϵ* and *ϵ*, and it has the same *d* dimensions as *x*. PGD is typically known as the most potent first-order attack, and this is due to the projection function.

#### 3.2.4. Carlini and Wagner (CW) Attack

The CW attack [14] is an advanced optimization-based attack. There are two types of the CW attack: *L*_2_ and *L*_∞_. We concentrate on the *L*∞ of targeted CW attack which can be solved by the PGD algorithm iteratively as following:

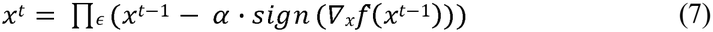

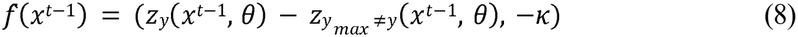

where *z*_*y*_ is the logits with respect to the class *y*, 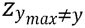 is the maximum logits of classes other than y, *f*(·) is the surrogate loss for the constraint restricted optimization problem stated in Eqn. (3), and *κ* is a factor that determines the attack confidence.

There also exist many more methods of adversarial attacks, in this paper, we attend to the four attacks mentioned above: FGSM, BIM, PGD and CW (figure 4).

**Figure 4.**
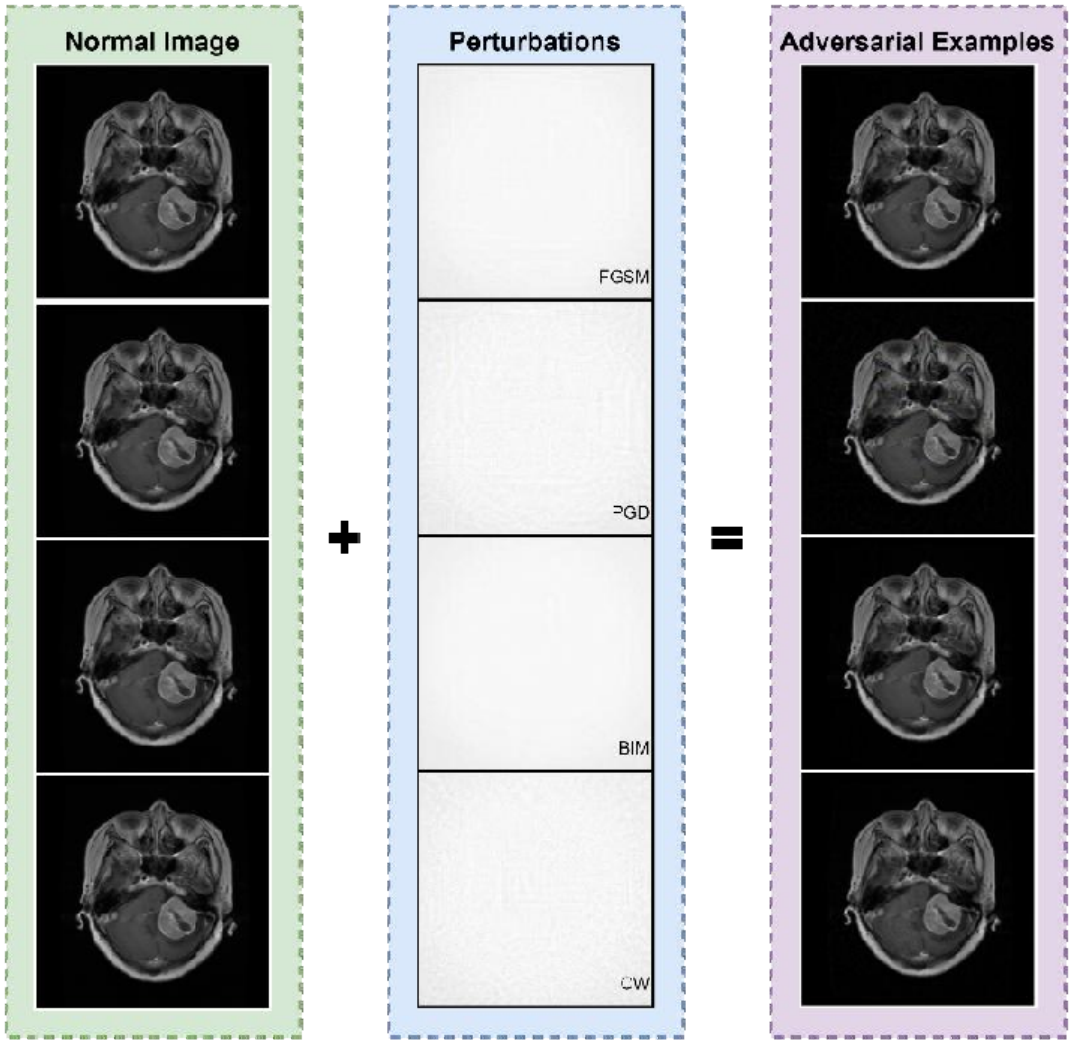
Examples of adversarial attacks crafted using FGSM, BIM, PGD and CW attack to fool Deep Neural Networks (DNNs) trained on brain MRI images. *Left:* normal images, *Middle:* adversarial perturbations (these panels may be magnified for discerning the various perturbations). *Right:* adversarial examples

## 4. RESULTS

### 4.1. Attack Settings

The attacks that we consider are: 1) FGSM, 2) BIM, 3) PGD, and 4) CW (_∞_ version). It should be noted that all these attacks are bounded attacks which means that, the maximum perturbation on each input pixel is not greater than a pre-defined maximum perturbation , with respect to _∞_ . These four types of attacks are produced on the test sets of the image dataset. To acquire the input gradients, based upon which the image is perturbed to maximize the network’s loss to the correct class, the input gradient extractor inputs the given image into the pre-trained DNN. For BIM, PGD and CW the perturbation steps are all set to 10, while the step size is set to */*10. As mentioned earlier, we focus on untargeted attacks in a white-box setting.

### 4.2. High susceptibility of neuroimaging scans to adversarial attack

In these sections, we explain the phenomenon observed above from two angles: 1) the properties of brain images, and 2) the properties of DNN models used.

#### 4.2.1. Medical Image Viewpoint

The attention map of an input image shows us the areas that interfere with the model output the most, based on the gradients of the classification loss with respect to the input. We can see that some medical images tend to have significantly larger regions of greater attention (figure 6). This could suggest that the DNN model can sometimes be distracted by the rich biological textures in brain images into paying more attention to regions that are not necessarily relevant to the diagnosis task. Even small perturbations in these attention regions can lead to major changes in the model output. In other words, this complexity of brain images makes it more vulnerable to attacks. However, this explanation does not provide the complete answer to the question, as we all know that even some natural images have a high level of complexity.

#### 4.2.2. DNN Model Viewpoint

Next, we illustrate that the increased susceptibility of DNN models for neuroimaging is majorly due to the use of overparameterized deep networks for basic neuroimage analysis tasks. To our surprise, we find that deep representations of brain imagery are very simple. This tells us that, DNN model is learning simple patterns from a large area of attention, which may only be lesion related. Learning simple patterns does not require sophisticated and over-complicated deep networks. This drives us to further examine about whether the high susceptibility is due to the use of overparameterized networks, which is done through exploring the loss landscape surrounding each input samples. Following up on earlier research on natural adversarial images [11, 13], we build adversarial direction g, where g is the input gradients taken from the DNN segmentation models and a set of surrogate models trained separate, respectively. We now create adversarial examples following the formula xadv = x +tab *ϵ*_1_g. To be more specific, we progressively increase *ϵ*_1_ from 0.01 to 0.1, and visualize the DICE score for *ϵ*_1_ in eqn.4. The loss landscapes surrounding brain images appear to be extremely clear. High susceptibility to adversarial attacks is a direct result caused by sharp loss; this is because small perturbations of an input sample make it likely to lead to a drastic increase in loss. Over complicated networks for a simple segmentation problem can cause sharp loss.

In summary, we have observed that DNN models for neuroimaging are susceptible to adversarial attacks, due to 2 reasons: 1) the intricate biological textures of brain images may result in areas that are more vulnerable; and most importantly, 2) Advanced deep networks made for processing large-scale natural images can turn out to be overparameterized for neuroimaging tasks making them highly susceptible to adversarial attacks.

### 4.3 CNN and ViT perform equally well on neuroimaging segmentation task

We then trained various CNNs (UNet, UNet++, DynamicUNet, DenseUNet, ResUNet) and ViTs (SwinUNet, TransUNet) on this task on brain tumor dataset. The resulting segmentation models performed well on the clean test set. ViT reached comparable baseline DICE scores of 0.43 for SwinUNet and 0.46 for TransUNet (Table 2), which were on par with CNN models such as UNet (0.437), DynamicUNet (0.435), and ResUNet (0.467) (Table 1).

**Table 1.**
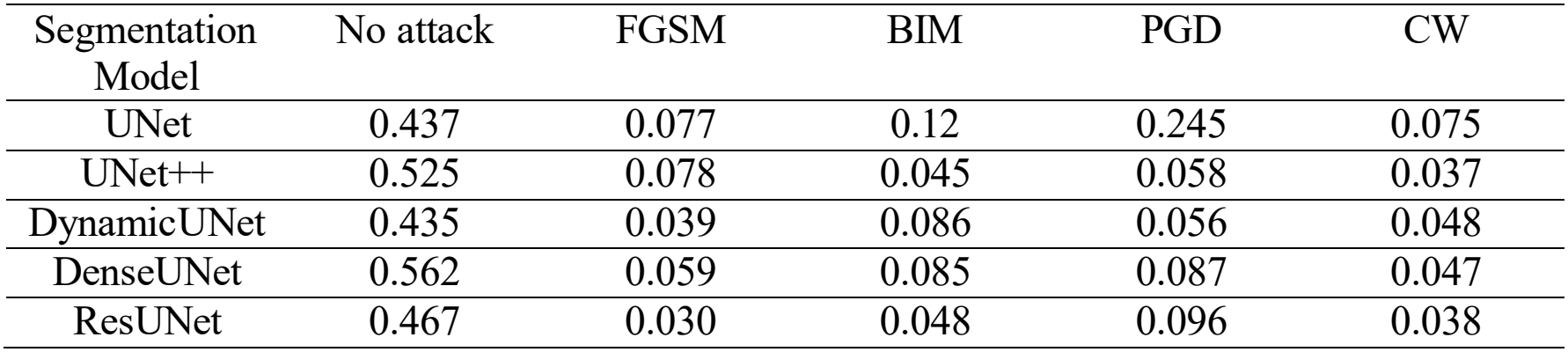
The DICE score of CNN based segmentation models on clean test images (denoted as “No attack”) and the 4 types of adversarial examples under *L*_∞_ maximum perturbation 0.01 for FGSM, BIM, PGD and CW attack.

| Segmentation Model | No attack | FGSM | BIM | PGD | CW |
|---|---|---|---|---|---|
| UNet | 0.437 | 0.077 | 0.12 | 0.245 | 0.075 |
| UNet++ | 0.525 | 0.078 | 0.045 | 0.058 | 0.037 |
| DynamicUNet | 0.435 | 0.039 | 0.086 | 0.056 | 0.048 |
| DenseUNet | 0.562 | 0.059 | 0.085 | 0.087 | 0.047 |
| ResUNet | 0.467 | 0.030 | 0.048 | 0.096 | 0.038 |

**Table 2.** The DICE score of ViT based segmentation models on clean test images (denoted as “No attack”) and the 4 types of adversarial examples under maximum perturbation 0.1 for FGSM, BIM, PGD and CW attack.

| Segmentation Model | No attack | FGSM | BIM | PGD | CW |
|---|---|---|---|---|---|
| SwinUNet | 0.43 | 0.28 | 0.17 | 0.19 | 0.29 |
| TransUNet | 0.46 | 0.17 | 0.16 | 0.27 | 0.26 |

### 4.4 CNNs are susceptible to multiple adversarial attacks

We now employed various adversarial attacks on CNNs, and evaluated white-box attacks. We used the most common and prevalent gradient-based attacks: Fast Gradient Sign Method (FGSM), Basic Iterative Method (BIM), Projected Gradient Descent (PGD), and Carlini and Wagner (CW) attack. We discovered that under adversarial perturbations (*ϵ* = 0.01), the segmentation performance of CNNs on the test set sharply decreased across all models (Table 1).

Specifically, for UNet, the DICE score dropped from a baseline of 0.437 to 0.077 under FGSM, 0.120 under BIM, 0.245 under PGD, and 0.075 under CW attack. Similarly, DenseUNet performance dropped from a baseline of 0.562 to 0.059 under FGSM and 0.047 under CW, while UNet++ dropped from 0.525 to 0.037 under CW. Across all CNN models, adversarial attacks severely degraded the segmentation performance. Taking into consideration all this data shows us that UNets and CNN-based models are highly vulnerable to adversarial attacks in neuroimaging.

### 4.5. Vision Transformers are inherently robust to adversarial attacks

Next, we targeted ViTs with various adversarial attacks. We discovered that even without any modification to the architecture they were relatively more resilient to adversarial attacks (Table 2). For SwinUNet under FGSM, BIM, PGD, and CW attacks (under maximum perturbation *ϵ* = 0.1), the DICE score retained values of 0.28, 0.17, 0.19, and 0.29 respectively, compared to its baseline of 0.43. Similarly, TransUNet maintained DICE scores of 0.17 (FGSM), 0.16 (BIM), 0.27 (PGD), and 0.26 (CW) from a baseline of 0.46.

Although evaluated under a higher perturbation parameter (*ϵ*= 0.1), ViTs preserved higher segmentation performance relative to their baseline compared to CNNs. Taking into consideration all this data shows us that ViTs are substantially more resilient to adversarial attacks than standard CNN architectures in neuroimaging.

### 4.6. Mechanism of robustness of Vision Transformers against adversarial attacks

To identify the reasons for this higher robustness exhibited by ViTs in response to adversarial attacks, we examined the adversarial noise obtained with white-box attacks on ViTs and UNets. We discovered that, quantitatively the magnitude of the gradients was consistently lesser for ViTs than for UNets (Table 2). Qualitatively, we could see that ViTs produced a distinct patch partition boundary alignment, and in the case of UNet patterns were more spatially inconsistent. This observation echoes the patch-based nature of ViTs, which acts to enable learned features to contain less low-level information such as lines and edges from an input image, which in turn makes them less sensitive to high-frequency perturbations.

Furthermore, applying dimensionality reduction via principal component analysis (PCA), the theoretical workflow of which is schematically outlined in Figure 5, we examined the structure of the latent space of the deep layer activations in UNets and ViTs. In case of the original images in the segmentation task, instances in the classes were more clearly visually separated for ViTs than for UNets. Quantitatively, and the distance between class centers were larger and the examples within a given class were aggregated more closely in the ViT latent space. When we attacked the images and used the baseline model to extract the features, the effects were even more pronounced: the UNet latent space was more de-clustered than the ViT latent space.

**Figure 5.**
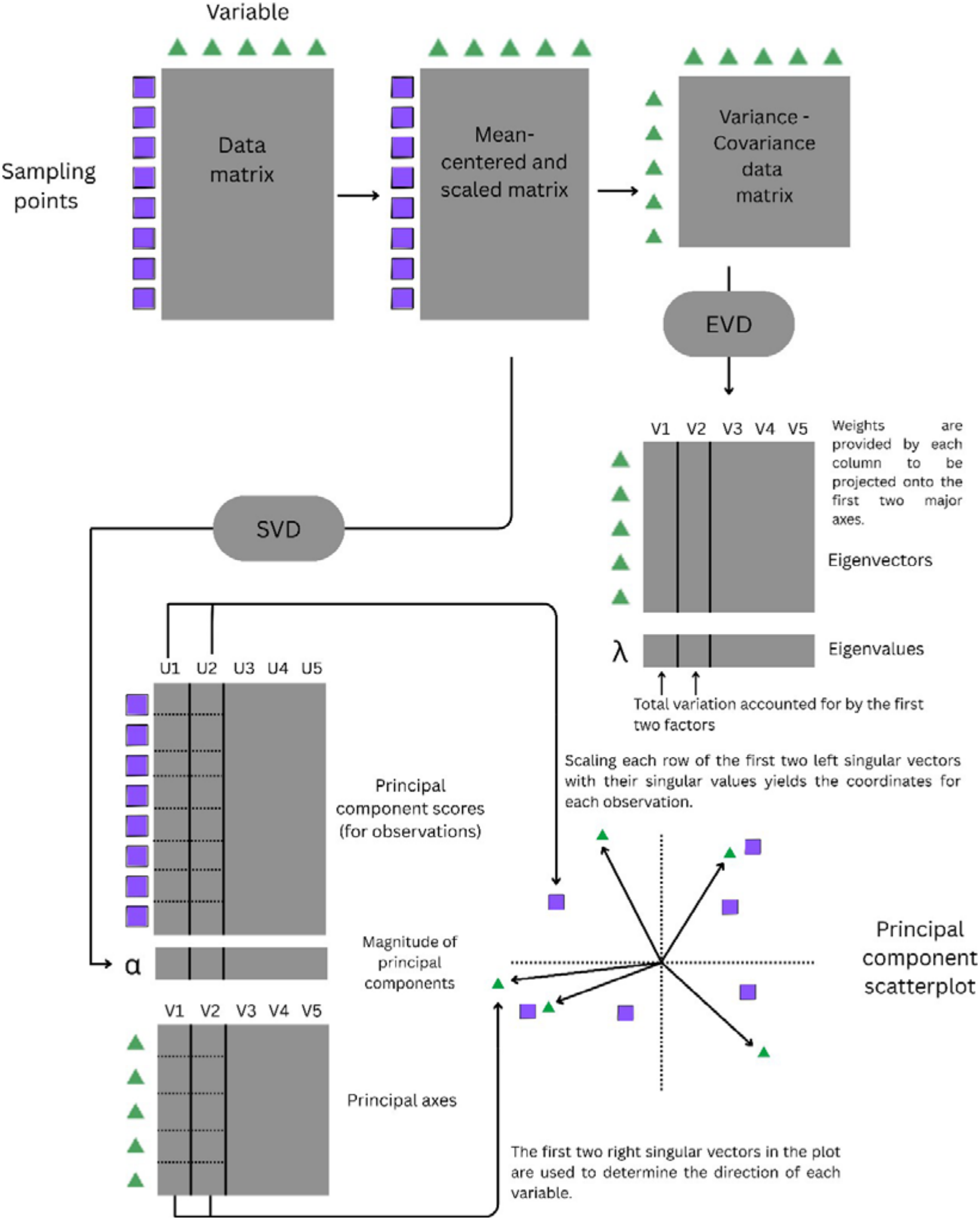
| Schematic representation of the PCA workflow.

**Figure 6.**
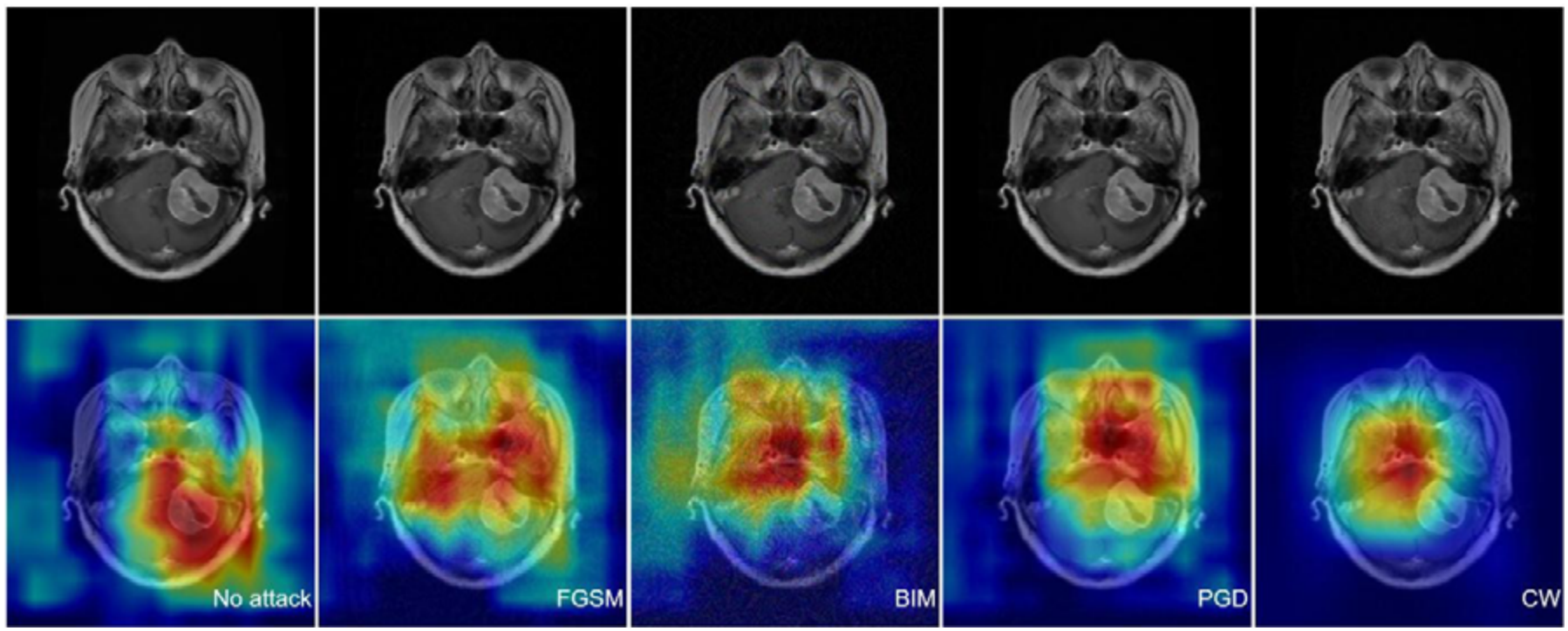
Attention maps of UNet-based Convolutional Neural Networks (CNNs) on normal brain MRI images versus adversarial images. These attention maps are generated using Grad-CAM.

Finally, utilizing attention maps we examined which regions in input images were given high importance by UNets and ViTs, respectively, using Grad-CAM [1] for visualization (figure 6 and figure 7). At baseline, the UNet would concentrate on a single region of the input image, while the ViT had multiple high attention image regions. After an adversarial attack, the importance of the UNet region became defocused and included much larger, potentially irrelevant image regions, with this effect increasing with attack strength (□). During an attack, there was no discernible change in the key image regions that Grad-CAM indicated in a ViT.

**Figure 7.**
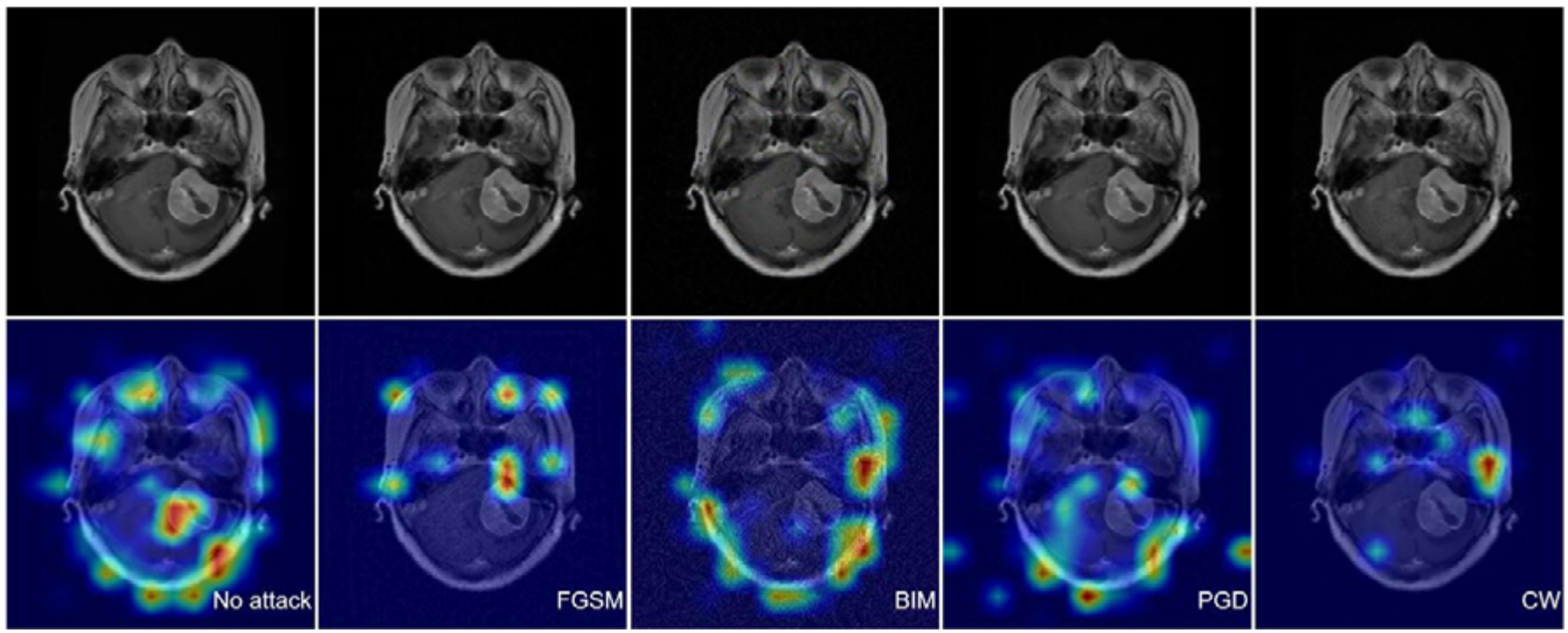
Attention maps of UNet-based Vision Transformers (ViTs) on normal brain MRI images versus adversarial images. These attention maps are generated using Grad-CAM.

In light of these findings, we conclude that the greater robustness of ViTs to white-box adversarial attacks, compared to UNets, is attributed to a more stable focus on relevant image regions within image tiles and its better separation of distinct classes in the latent space.

## 5 DISCUSSION

Machine learning (ML) based software as medical devices (SaMD) can be a target of cyberattacks, which have the potential to seriously disrupt the medical diagnosis. AI systems can be manipulated by adversarial attacks into making incorrect and inaccurate predictions. In the health care field, the Use of AI systems is ever growing. One specific, relevant domain of application is neuroimaging, where AI systems have been used to answer several medically important questions. These academic advancements have led to the development of many advanced AI algorithms. Given the information that is publicly available, these algorithms are presumably mostly based on CNNs, rather than ViTs. In the end, diagnostic stakeholders may benefit from these algorithms’ potential for efficiency and resource savings. However, it’s crucial to make sure AI systems are resilient to hostile manipulations and artifacts throughout any possible large-scale deployment.

Here, we demonstrate that CNNs used for neuroimaging tasks are susceptible to adversarial attacks well below the threshold for human perception. We investigate varied and commonly used UNet models, and prove that all are equally vulnerable to adversarial attacks. To solve this issue, we investigated the potential of ViT models to exhibit adversarial robustness. We show that ViTs perform similar to UNets at baseline, and that they seem to be inherently more robust against adversarial attacks. Additionally, we also noticed that the more complicated models with a greater number of trainable parameters are more susceptible to adversarial attacks, but ViT proves to be robust despite its large number of parameters. Despite the fact that no AI models are universally and fully impervious to attacks, our research and its results demonstrates that ViTs are much more robust against common white-box attack types and that this is attributed to a more robust behaviour of the latent space compared to CNNs. Our results support the use of ViTs as the foundational technology for AI products in neuroimaging and add to a list of theoretical advantages of ViTs over CNNs.

## 6 CONCLUSIONS

In neuroimaging diagnostics, it is a common practice to use advanced deep neural networks developed initially for complex, large-scale natural image processing. However, these networks when used for brain imaging tasks, results in a sharp loss of performance brought about by the use of an overcomplicated network on lesion or tissue segmentation task. We would like to highlight that these diagnostic systems are more vulnerable to adversarial attacks. We have also shown that the contemporary AI approach of Vision Transformers furnishes enhanced structural resilience to automated brain lesion segmentation procedure in neuroimaging during hostile digital intrusion. Our investigations imply that commercial adoption of AI neuroimaging diagnostics, should rely on the Vision Transformer methodology, instead of the customary CNN technique, for segmentation of pathological regions of interest; this has a novel potentiality for practical impact as there can be enhanced diagnostic protection against malicious digital hostility.

### Credit authorship contribution statement

#### Mayur Mankar

Conceptualization, Data Curation, Formal analysis, Investigation, Methodology, Software, Validation, Visualization, Writing – original draft, Writing – review and editing. *Prasun K. Roy:* Conceptualization, Formal analysis, Investigation, Methodology, Validation, Project Administration, Visualization, Supervision, Writing – original draft, Writing – review and editing.

### Funding sources

The authors PKR and MM greatly appreciate the support from C3i Hub Foundation (sponsored by DST: Department of Science and Technology, Govt. of India).

MM appreciates the support of travel fellowship from Indian Academy of Sciences, Bangalore.

## Acknowledgement

We acknowledge the help of La Dassault Systems Foundation, France, for logistic support.

## REFERENCES

[1] Selvaraju, R., et al, “Grad-CAM: Visual Explanations from Deep Networks via Gradient-Based Localization,” in 2017 IEEE International Conference on Computer Vision (ICCV), 2017, pp. 618–626

[2] Cheng, Q. (2015). Enhanced Performance of Brain Tumor Classification via Tumor Region Augmentation and Partition. PLOS ONE, 10(10), 1–13. Link of the scansets: 10.6084/m9.figshare.1512427

[3] Olaf Ronneberger, Philipp Fischer, & Thomas Brox. (2015). U-Net: Convolutional Networks for Biomedical Image Segmentation.

[4] Zongwei Zhou, Md Mahfuzur Rahman Siddiquee, Nima Tajbakhsh, & Jianming Liang. (2018). UNet++: A Nested U-Net Architecture for Medical Image Segmentation.

[5] Jin Yang, Daniel S. Marcus, & Aristeidis Sotiras. (2024). Dynamic U-Net: Adaptively Calibrate Features for Abdominal Multi-organ Segmentation.00020

[6] Cao, Y., Liu, S., Peng, Y., & Li, J. (2020). Denseunet: densely connected unet for electron microscopy image segmentation. IET Image Processing, 14(12), 2682–2689. 10.1049/iet-ipr.2019.1527

[7] Anita Khanna, Narendra D. Londhe, S. Gupta, & Ashish Semwal (2020). A deep Residual U-Net convolutional neural network for automated lung segmentation in computed tomography images. Biocybernetics and Biomedical Engineering, 40(3), 1314–1327.

[8] Dosovitskiy, A., Beyer, L., Kolesnikov, A., Weissenborn, D., Zhai, X., Unterthiner, T., Dehghani, M., Minderer, M., Heigold, G., Gelly, S., Uszkoreit, J., & Houlsby, N. (2020b). An Image is Worth 16x16 Words: Trans11formers for Image Recognition at Scale. arXiv (Cornell University). 10.48550/arxiv.2010.11929

[9] Cao, M. (2023). Swin-Unet: Unet-Like Pure Transformer for Medical Image Segmentation. In Computer Vision – ECCV 2022 Workshops (pp. 205–218). Springer Nature Switzerland.

[10] Jieneng Chen, Yongyi Lu, Qihang Yu, Xiangde Luo, Ehsan Adeli, Yan Wang, L. Lu, Alan L. Yuille, & Yuyin Zhou. (2021). TransUNet: Transformers Make Strong Encoders for Medical Image Segmentation.

[11] Goodfellow, I. J., Shlens, J., & Szegedy, C. (2015). Explaining and Harnessing Adversarial Examples. International Conference on Learning Representations (ICLR).

[12] Kurakin, A., Goodfellow, I., & Bengio, S. (2016). Adversarial examples in the physical world. arXiv preprint arXiv:1607.02533.

[13] Madry, A., Makelov, A., Schmidt, L., Tsipras, D., & Vladu, A. (2018). Towards Deep Learning Models Resistant to Adversarial Attacks. International Conference on Learning Representations (ICLR).

[14] Carlini, N., & Wagner, D. (2017). Towards Evaluating the Robustness of Neural Networks. 2017 IEEE Symposium on Security and Privacy (SP), 39–57.

